# 14-3-3ζ-depletion impairs mammary gland development in the mouse

**DOI:** 10.1101/2021.02.03.425513

**Authors:** Valentina Poltavets, Zahra Esmaeili, Sarah T. Boyle, Hayley S. Ramshaw, Angel F. Lopez, Marina Kochetkova, Michael S. Samuel

## Abstract

The 14-3-3 family of proteins have roles in regulating several key cellular processes. While their significant structural and functional homology had informed the idea that these proteins acted redundantly, it is now becoming clear that individual family members may have tissue and context specific functions, highlighting the need for a more nuanced understanding of these important proteins. Here, we demonstrate that mice deficient in 14-3-3ζ exhibit developmental defects of the mammary epithelium, associated with dysregulation of key transcription factors involved in the maintenance of mammary stem cell populations. We believe that this model will be prove useful for investigating the role of 14-3-3ζ in the maintenance of mammary stem cell populations and elucidating the transcriptional networks driving specification of the mammary epithelium.

## Introduction

The mammary gland is an organ unique to mammals, with the specific function of milk production and delivery. It develops in a series of three defined stages – embryonic, pubertal and reproductive, with each stage initiated in response to specific biochemical and biomechanical cues (Anlas & Nelson, 2018; Macias & Hinck, 2012). Pubertal development of the mammary gland in mice is initiated at around 4 weeks of age and occurs via a process of branching morphogenesis of a pre-existing rudimentary ductal network, driven by the proliferation of terminal end buds (TEBs), which results in elongation of the mammary ducts. Lateral branching from primary ducts leads to the elaboration of a tree-like structure that constitutes the mammary epithelium. This process is orchestrated by hormonal signals from the ovary and the pituitary (Topper & Freeman, 1980). While key differences exist, the mouse mammary gland remains an excellent model to study the development of this organ in humans, with each stage of human breast development mirrored by analogous processes in the mouse, and indeed significant scientific insight into human development has already been gained by detailed study of this model (McNally & Stein, 2017).

14-3-3ζ, encoded by the *Ywhaz* gene, is a member of a family of phospho-serine binding proteins (with other family members designated β, γ, ε, σ, τ and η) that are highly conserved across both the plant and animal kingdoms, where they regulate diverse cellular processes by acting as molecular chaperones and regulating protein activity, stability and sub-cellular localization, acting as obligate dimers to engage with phosphorylated Ser/Thr residues (Camoni, Visconti, Aducci, & Marra, 2018). Little is known about the functions of 14-3-3 family members in normal mammary gland development, but it appears that several family members function non-redundantly in this process. For example, 14-3-3σ was found to be important in maintaining epithelial polarity in normal murine mammary cells (Ling, Zuo, Xue, Muthuswamy, & Muller, 2010) and 14-3-3ε has been reported to regulate cell proliferation and casein synthesis via PI3K-mTOR signaling in bovine mammary epithelial cells (Wang, Wang, Yang, Liu, & Jiang, 2018). A further study demonstrated, also in bovine mammary epithelial cells, that 14-3-3γ regulates protein synthesis and cell proliferation at both translational and transcriptional levels and that mTOR, Stat5a and β-casein were positively regulated by 14-3-3γ overexpression (Yu et al., 2014). While these findings show that the 14-3-3 family members are important in mammary development, they also suggest that individual 14-3-3 family members perform non-redundant functions in this process.

The 14-3-3 proteins are the subject of intense research as mediators of protein-protein interactions and therefore potentially exploitable as mimetics or targets for a variety of pathological conditions (Stevers et al., 2018). We have previously reported the rational design and validation of a family of pharmacological inhibitors of 14-3-3 proteins that function by promoting phosphorylation at Ser-58, located at the dimer interface, thereby preventing the formation of 14-3-3 dimers (Woodcock et al., 2015). However, before these can be tested for therapeutic efficiency, a comprehensive understanding of their biology, particularly their roles in mammalian development, is essential to circumvent deleterious off-target effects. It has therefore been necessary to generate and validate gene-targeted mice lacking individual 14-3-3 family members. Here, we characterize mammary development in mice lacking 14-3-3ζ, which we propose would be a useful model to understand the role of this important molecular chaperone in regulating developmental processes at both the local level and via systemic changes.

## Results and Discussion

The generation of mice bearing the targeted insertion of a gene trap into the first intron of *Ywhaz* from the Lexicon Genetics ES cell line OST062 has been previously described (Cheah et al., 2012). To establish that this gene trap depletes *Ywhaz* transcript in mammary glands, we undertook quantitative real-time polymerase chain reaction (qPCR) analysis of cDNA synthesized from mRNA derived from mammary duct preparations, using a primer pair that is specific for *Ywhaz*. These analyses revealed that there is no detectable *Ywhaz* transcript in homozygous *Ywhaz* gene trap mice (KO), whereas mice that were heterozygous for the *Ywhaz* gene trap (HET) exhibited half the level observed in wild-type (WT) mice (Figure 1A). Accordingly, quantitative immunofluorescence analysis of pubertal mammary fat pads (MFP) using an antibody that selectively binds to 14-3-3ζ protein yielded no signal in KO MFPs, but around half the level of fluorescence signal in HET MFPs compared to WT MFPs (Figure 1B). These data suggest that this gene targeting approach abolished expression of 14-3-3ζ in the MFP at both the transcript and protein levels and that heterozygous mice express approximately half the levels of transcript and protein in their MFPs.

**Figure 1.**
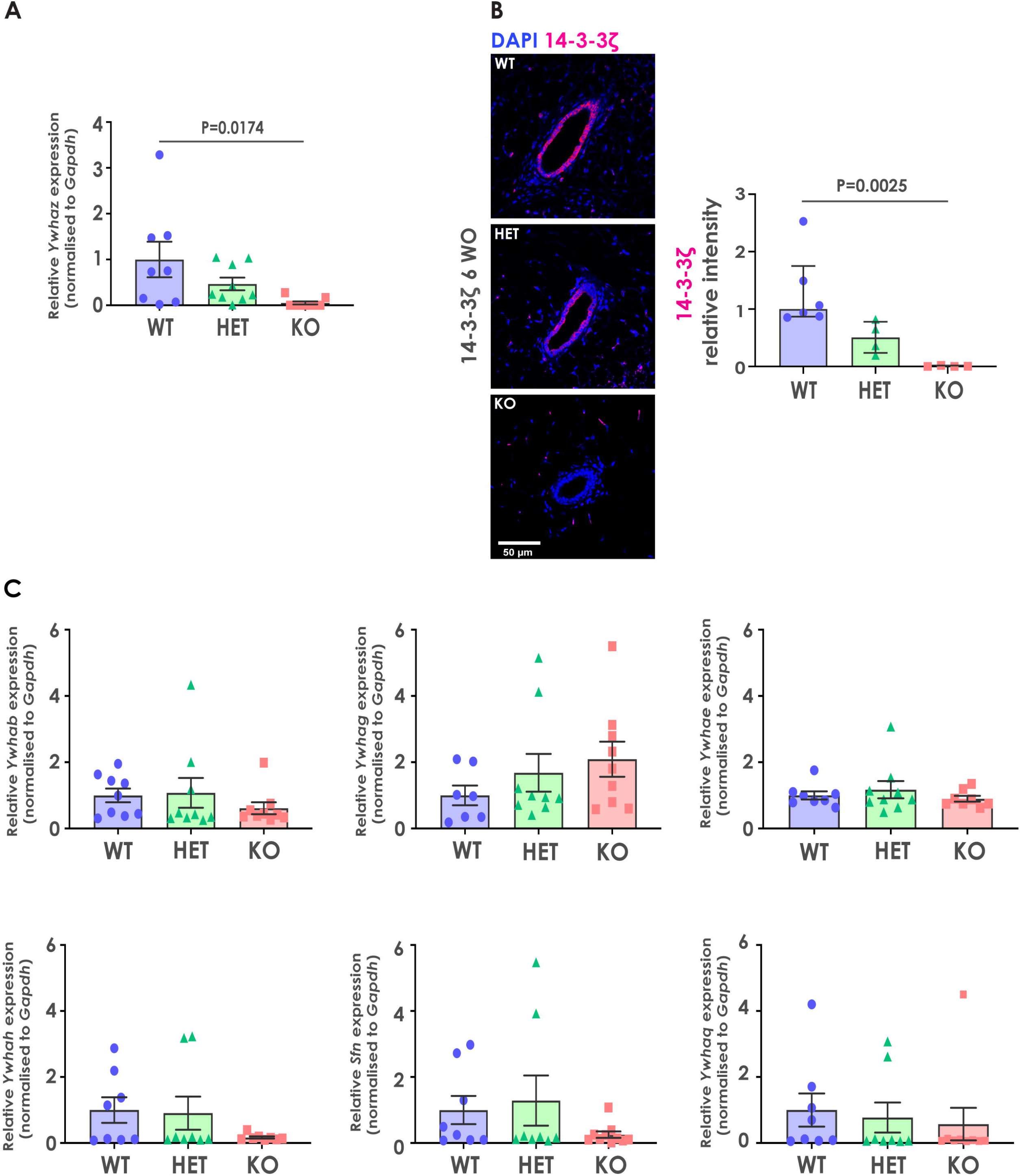
(A) *Ywhaz* gene expression analysis of the mammary epithelia of 14-3-3ζ wild-type (WT), heterozygous (HET) and deficient (KO) mice, expressed relative to *Gapdh* and shown as Mean ± standard error of the mean (SEM). P values determined by one-way ANOVA. N=8(WT) and 9(HET, KO) individual mammary fat pads. (B) Representative images of immunofluorescence analysis of 14-3-3ζ in WT, HET and KO mammary glands of 6-week-old mice. Signal intensity was quantified per area of ductal epithelium and expressed relative to WT. Data are shown as medians ± inter-quartile ranges (IQRs). P values were determined using the Kruskal–Wallis test. N=6 (WT), 4 (HET, KO) individual mammary fat pads. (C) Gene expression analysis of genes encoding 14-3-3 family members as indicated, expressed relative to *Gapdh* and shown as medians ± IQRs. Each dot indicates a separate mammary fat pad.

We next assessed whether the other six 14-3-3 family members were differentially regulated in KO and HET mice relative to WT littermates. To this end, we conducted qPCR analyses on RNA isolated from mammary duct preparations from WT, HET and KO mice, using primer pairs specific for *Ywhab* (which encodes 14-3-3β), *Ywhae* (encoding 14-3-3ε), *Ywhag* (encoding 14-3-3γ), *Ywhah* (encoding 14-3-3η), *Ywhaq* (encoding 14-3-3τ) and *Sfn* (encoding 14-3-3σ) transcripts. These analyses revealed that there is no significant differential regulation of the other 14-3-3 family members when 14-3-3ζ is depleted within the mammary gland (Figure 1C).

We next sought to systematically assess mammary development in 14-3-3ζ-deficient mice relative to HET and WT mice. Gross morphology of the inguinal (4^th^) mammary glands was compared in WT, HET and KO mice on the FVB/n background at 4, 6, 9 and 11 weeks of age. These ages capture key stages of mammary development in mice of this strain background, namely onset and early puberty (3-6 weeks), mid-puberty (7-10 weeks) and maturity (∼10-12 weeks) (Richert, Schwertfeger, Ryder, & Anderson, 2000). Although the mammary glands of KO mice appeared identical to those of WT mice at 4 weeks of age, we observed a pronounced developmental delay in epithelial branching at 6 and 9 weeks of age in KO animals, which persisted till 11 weeks of age (Figure 2A). However, the mammary glands of the WT and HET animals were developmentally indistinguishable throughout this age range (Figure 2A).

**Figure 2.**
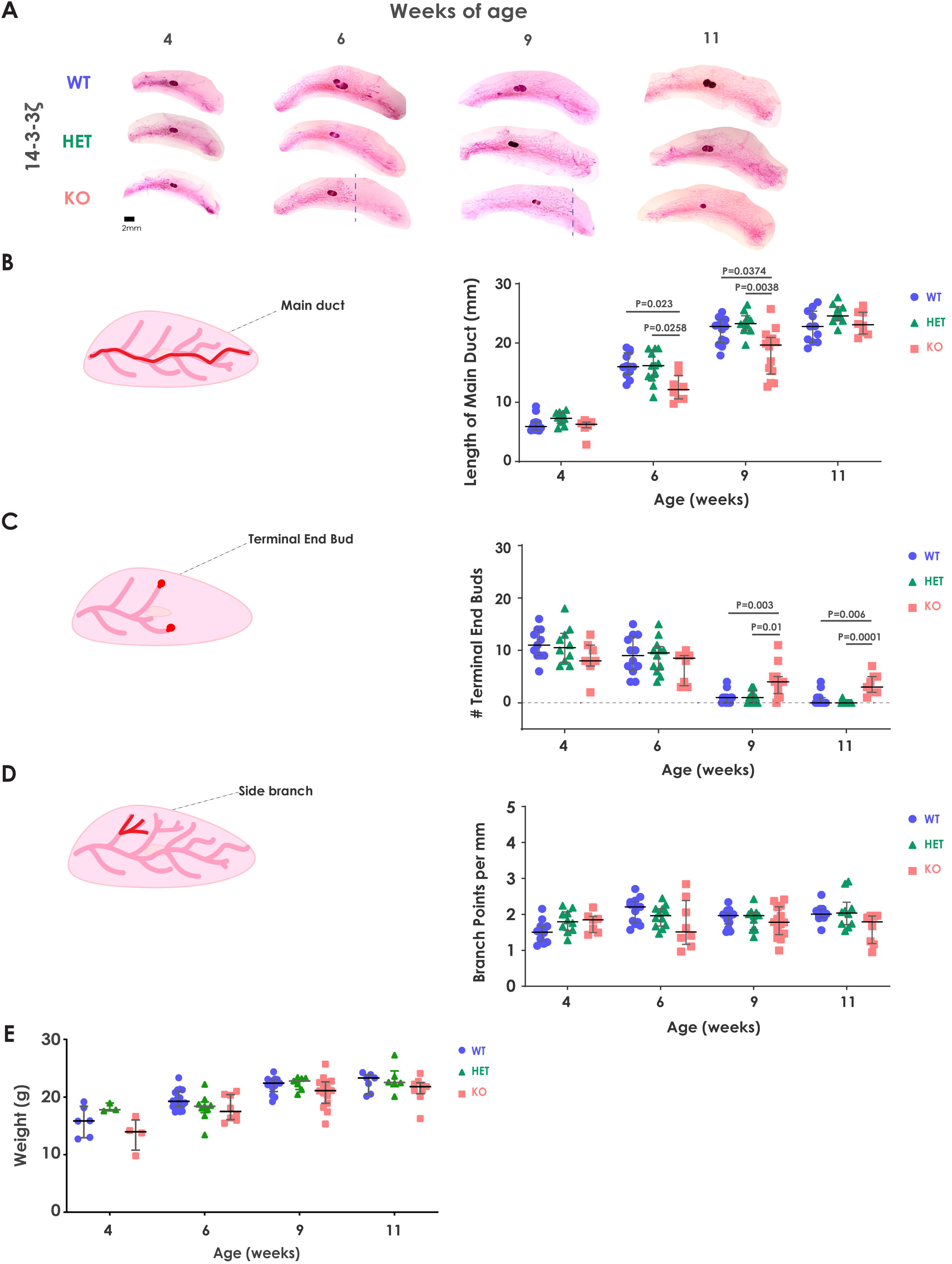
(A) Representative images of whole mount 4th mammary glands taken from WT, HET and KO mice at 4-11 weeks of age. The extent of mammary tree elaboration of KO animals is indicated by the dotted line. (B) Quantification of the length of the main mammary epithelial duct from WT, HET and KO mice at 4-11 weeks of age. Data are shown as medians ± IQRs, compared using the Kruskal-Wallis test (N=7-14 MFPs per genotype). Each dot represents a separate mammary fat pad. (C) The number of the terminal end buds (TEBs) in WT, HET and KO mice at 4-11 weeks of age. TEBs are defined as teardrop-shaped structures larger than 0.03 mm^2^ in area at the end of the duct. Data shown as medians ± IQRs, analyzed using the Kruskal-Wallis test (N=7-14 MFPs per genotype). Each dot indicates a separate mammary fat pad. (D) Quantification of the secondary branching within mammary epithelium determined by measurement of branch points per mm along three separate ducts. Data are shown as medians ± IQRs, analyzed using the Kruskal-Wallis test (N=7-14 MFPs per genotype). Each dot indicates a separate mammary fat pad. (E) Body weights of WT, HET and 14-3-3ζ-deficient mice at 4-11 weeks of age used in the above analyses. Data shown as medians ± IQRs. Each dot indicates a separate mouse (N=3-17 mice).

The rudimentary epithelial structures of the murine mammary gland remain quiescent until ∼3 weeks of age when the onset of puberty, triggered by sex hormones, leads to the appearance of terminal end buds (TEBs) -the main proliferative centers of the mammary tree, which are enriched for stem cells and undifferentiated progenitor cells relative to mature ducts. The process of ductal elongation ensues and continues till ∼8-9 weeks of age. At ∼10-12 weeks the TEBs reach the physical limits of the fat pad and differentiate into developed mammary ducts, completing the process of mammary epithelial elaboration (Richert et al., 2000). To examine the process of branching morphogenesis in more detail, we characterized mammary epithelial outgrowth using several well-accepted quantification approaches that are informative of mammary development in vivo (Blacher et al., 2016). The duct beginning at the nipple and extending furthest of all ducts along the MFP at any point in development is termed the main duct (Schematic, Figure 2B). WT and HET MFPs exhibited similar main duct lengths at all timepoints examined and had completely traversed the MFP by 9 weeks. However, the main ducts of KO MFPs were 1.5-fold shorter than that of WT MFPs at weeks 6 and 9, and only reached their full length at 11 weeks of age (Figure 2B). WT and HET MFPs exhibited similar numbers of TEBs at each developmental stage, and these had disappeared by 9 weeks of age, consistent with the establishment of a mature mammary epithelial tree. KO MFPs, on the other hand, exhibited 3.5-fold more TEBs than WT and HET MFPs at both 9 and 11 weeks of age, indicative of a developmental delay (Figure 2C). Interestingly, TEBs persisted in 11-week-old KO mice even though elaboration of the mammary ductal structures in these mice was complete, further highlighting the developmental delay. To assess secondary branching during mammary ductal elaboration, we enumerated branchpoints per mm of duct length in all genotypes. Secondary branching in WT and HET MFPs was very similar, whereas KO MFPs exhibited a trend towards fewer branch points at 6, 9 and 11 weeks of age (Figure 2D).

Since the body weight of the animal could be a potential co-variate and thereby influence the evaluation of mammary gland development (Blacher et al., 2016), we weighed each animal prior to dissecting out MFPs. Bodyweights of mice of all genotypes were comparable at each timepoint (Figure 2E) and therefore, the delayed mammary development observed in KO mice relative to their WT and HET littermates cannot be explained by differences in overall body weight.

14-3-3 protein family members have established roles in cell proliferation, survival and differentiation (Mhawech, 2005). We therefore investigated whether 14-3-3ζ deficiency influenced cell proliferation in the developing mammary epithelium. At 6 weeks of age, the time at which we observed the biggest differences in mammary epithelial elaboration between WT and KO mice, immunohistochemical analysis revealed 15% fewer Ki-67 positive cells in the KO mammary epithelium relative to WT epithelium, whereas the number of Ki-67 positive cells in the mammary epithelium in HET animals was similar to that in WT mice (Figure 3A). We next investigated rates of cell death in the mammary epithelia of mice of all three genotypes. Terminal deoxynucleotidyl transferase dUTP nick end labeling (TUNEL) analysis revealed no discernable differences in cell death within the mammary epithelia of 6-week-old WT, HET or KO mice (Figure 3B). Taken together, these data suggest that impaired cell proliferation may contribute to impaired mammary ductal elaboration in KO mice. The mechanism by which 14-3-3ζ regulates cell proliferation within the MFP could be caused either by systemic effects of 14-3-3ζ deficiency resulting in aberrant hormone regulation or local effects within the epithelial and/or stromal components of the MFP.

**Figure 3.**
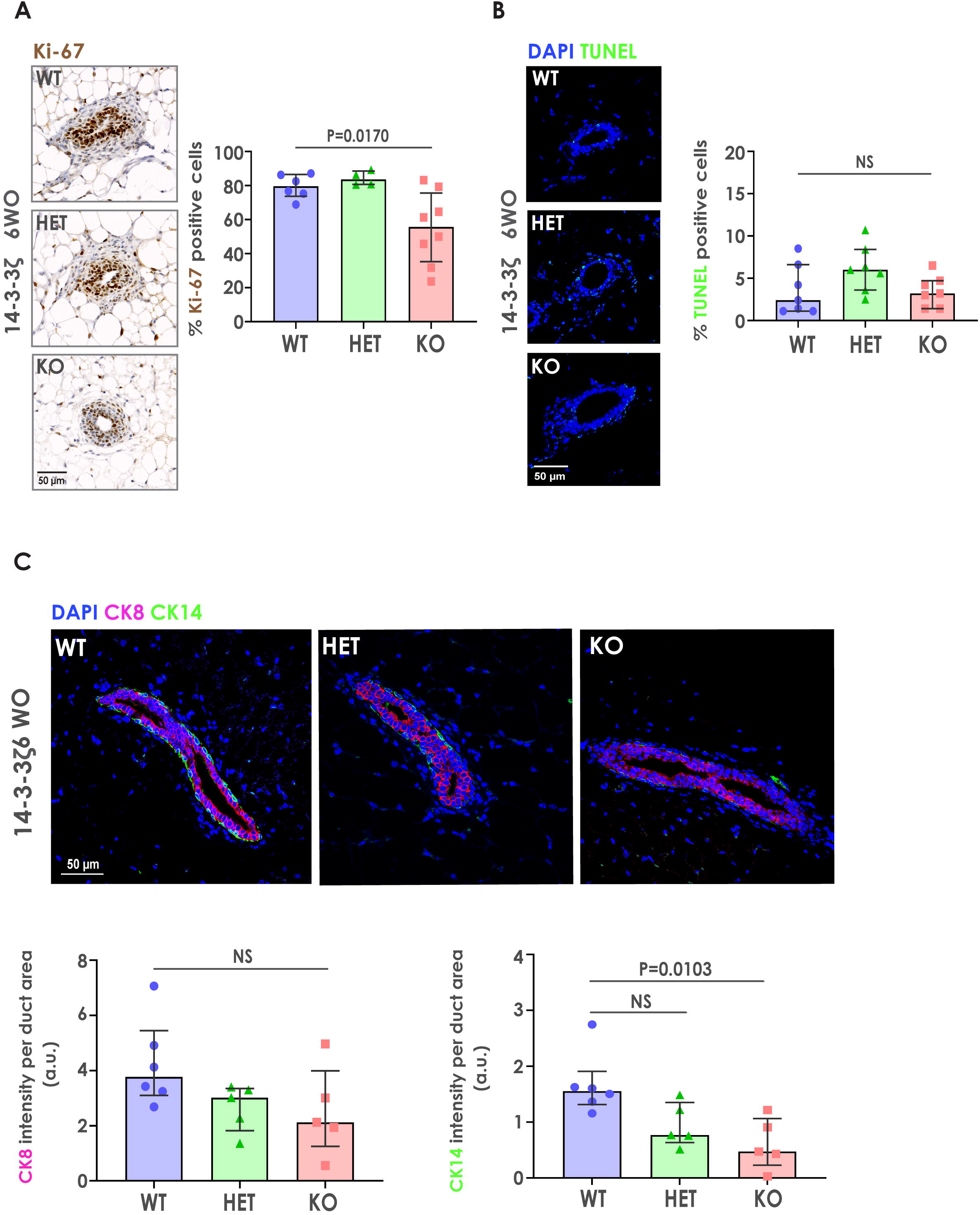
(A) Representative images and quantification of Ki-67 immunohistochemical staining in the mammary glands of 14-3-3ζ WT, HET and KO mice at 6 weeks of age. Data are shown as medians ± IQRs with P values determined by one-way ANOVA, (N=6 (WT), 4 (HET), 8 (KO) individual mammary fat pads). (B) Representative images and quantification of cell death via the TUNEL assay, in the mammary glands of 14-3-3ζ WT, HET and KO mice at 6 weeks of age. Data are shown as medians ± IQRs with P values determined by one-way ANOVA, (N=7, individual mammary fat pads). Each dot indicates a separate mammary fat pad. (C) Representative images of cytokeratin 8 (luminal) and cytokeratin 14 (basal) immunofluorescence analysis of mammary ducts of 6-week-old mice. Data are shown as medians ± IQRs and analyzed using the Kruskal-Wallis test. N=6 (WT), 5 (HET), 5 (KO) individual mammary fat pads. Each dot indicates a separate mammary fat pad.

Production of cytokeratin 8 (CK8) or cytokeratin 14 (CK14) are used to distinguish between luminal and basal epithelial populations of the mammary ducts respectively (Mikaelian et al., 2006). Immunofluorescence analysis of the mammary glands of 6-week-old mice revealed no significant differences between CK8+ luminal cell populations, but a significant depletion of CK14+ basal cell population in the mammary epithelium of KO mice relative to those of WT and HET mice (Figure 3C). These data suggest defects in basal cell lineage specification in the mammary epithelium of mice deficient for 14-3-3ζ.

A recent study has revealed the role of the transcription factor Dachshund homolog 1 (*Dach1*) as a key regulator of branching morphogenesis of the mammary ducts and the specification of basal mammary epithelial cells via its role in the maintenance of the pool of mammary progenitor cells (Jiao et al., 2019). We therefore undertook comparative gene expression analysis for *Dach1* in mammary epithelial cell preparations of WT, HET and KO MFPs. These analyses revealed a ∼75% lower level of *Dach1* gene expression in KO mammary epithelium relative to that of WT and HET mice (Figure 4A). Our data are therefore consistent with the previously reported role for *Dach1* in mammary ductal branching and suggest that 14-3-3ζ has a function in regulating *Dach1* levels during mammary gland development. Dach1 has been demonstrated to directly regulate the transcription of *Notch1, Pygo2* and *Gata3*, genes known to regulate mammary development (Jiao et al., 2019). Consistent with this report, we observed significantly lower levels of *Notch1* and *Pygo2* in KO mammary epithelium relative to WT and HET mammary epithelium at 6 weeks and a similar trend in *Gata3* levels at the same time point (Figure 4B). Taken together, these data suggest that 14-3-3ζ has a novel role in regulating the transcription factor Dach1 within the developing mammary epithelium, with potential implications also for broader developmental processes. While it remains to be established whether this is a direct or an indirect role, it is worth noting that the murine Dach1 protein contains 14-3-3-binding motifs around phospho-serine residues at positions 499 and 508 as identified by Scansite 4.0 (Obenauer, Cantley, & Yaffe, 2003). Changes in the stability or function of Dach1 protein has the potential to influence the transcription of its target genes and to feed back on its own regulation.

**Figure 4.**
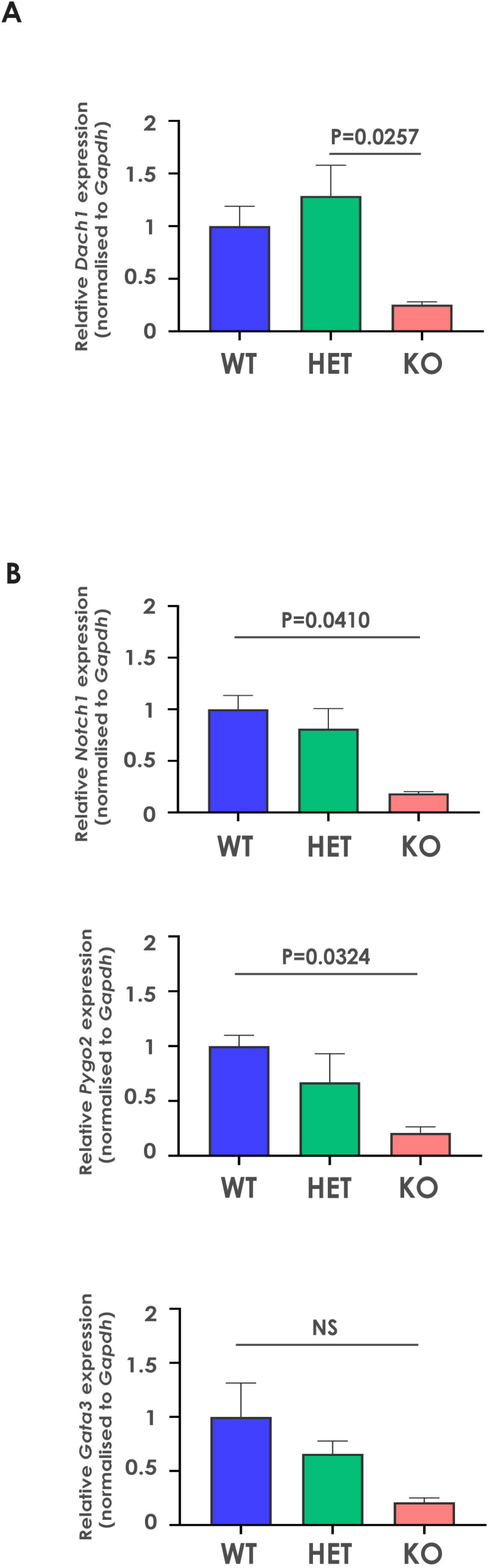
(A) Gene expression analysis of *Dach1*, in the mammary glands of 6-week-old WT, HET and KO mice relative to *Gapdh*. Data are relative to mean expression in WT mammary fat pads. Data are Mean ± SEM, analyzed using one-way ANOVA; N=3 mice. (B) Gene expression analysis of the Dach1 target genes, *Notch1, Pygo2* and *Gata3* in the mammary glands of 6-week-old WT, HET and KO mice relative to *Gapdh*. Data are relative to mean expression in WT mammary fat pads. Data are MeanL±LSEM, analyzed using one-way ANOVA; N=3 mice.

In conclusion, we report that mice selectively deficient for 14-3-3ζ exhibit disrupted mammary gland development associated, at least in part, with dysregulation of *Dach1* gene expression and that of its transcriptional targets. We propose that this model may be useful for investigating the role of 14-3-3ζ in the maintenance of mammary stem cell pools and the transcriptional network involved in specification of the mammary epithelium.

## Methods

### Mice

All procedures were approved by the institutional animal ethics committees of the Central Adelaide Local Health Network/SA Pathology and the University of South Australia. Generation of *Ywhaz* gene-targeted mice has been previously described (Cheah et al., 2012). Mice were originally generated on the 129/Sv strain background and subsequently backcrossed onto the FVB/n strain background for at least 10 generations to obtain 14-3-3ζ-deficient FVB/n mice.

### Mammary gland whole mount preparations, imaging and analysis

Whole mount preparations and quantification were carried out as previously described (Boyle et al., 2016). 4^th^ inguinal mammary fat pads were dissected, fixed in Carnoy’s fixative, then stained in Carmine Alum overnight at room temperature. Samples were then dehydrated through graduated ethanol, cleared in xylene and mounted using Permount (Thermo Fisher Scientific). Tissues were visualized using a dissecting microscope (Model MVX10, Olympus) connected to a CCL12 camera. Image stitching and processing was performed using MosaicJ plugin (http://bigwww.epfl.ch/thevenaz/mosaicj/) for ImageJ (NIH).

### Isolation of mammary epithelial cells

Total mammary cells were derived as previously described (Boyle et al., 2020). Briefly, all mouse mammary glands were dissected, manually dissociated and then digested in Dulbecco’s modified Eagle’s medium (DMEM, Gibco) supplemented with 1 mg/ml collagenase IA, 100U/ml hyaluronidase (Worthington Biochemical Corporation) and 2% fetal calf serum for 3 h at 37 °C. Resultant total cell preparations were processed in TRIzol® (Invitrogen) using The FastPrep-24® (MP Biomedicals) tissue homogenizer.

### Quantitative real-time polymerase chain reaction

RNA was extracted using TRIzol® according to the manufacturer’s instructions. RNA concentration was determined by NanoDrop spectrometer (Thermofisher). First strand cDNA synthesis was carried out using the QuantiTect Reverse Transcription Kit (Qiagen) according to the manufacturer’s instructions. Validated qPCR primers and primer assays (Qiagen, see Table 1) were used together with QuantiTect SYBR Green PCR kit (QIAGEN) for gene expression analysis on a Rotor-Gene Q(CA) thermocycler (QIAGEN). Data processing was performed using Rotor-Gene Q (CA) software (QIAGEN).

**Table 1:**
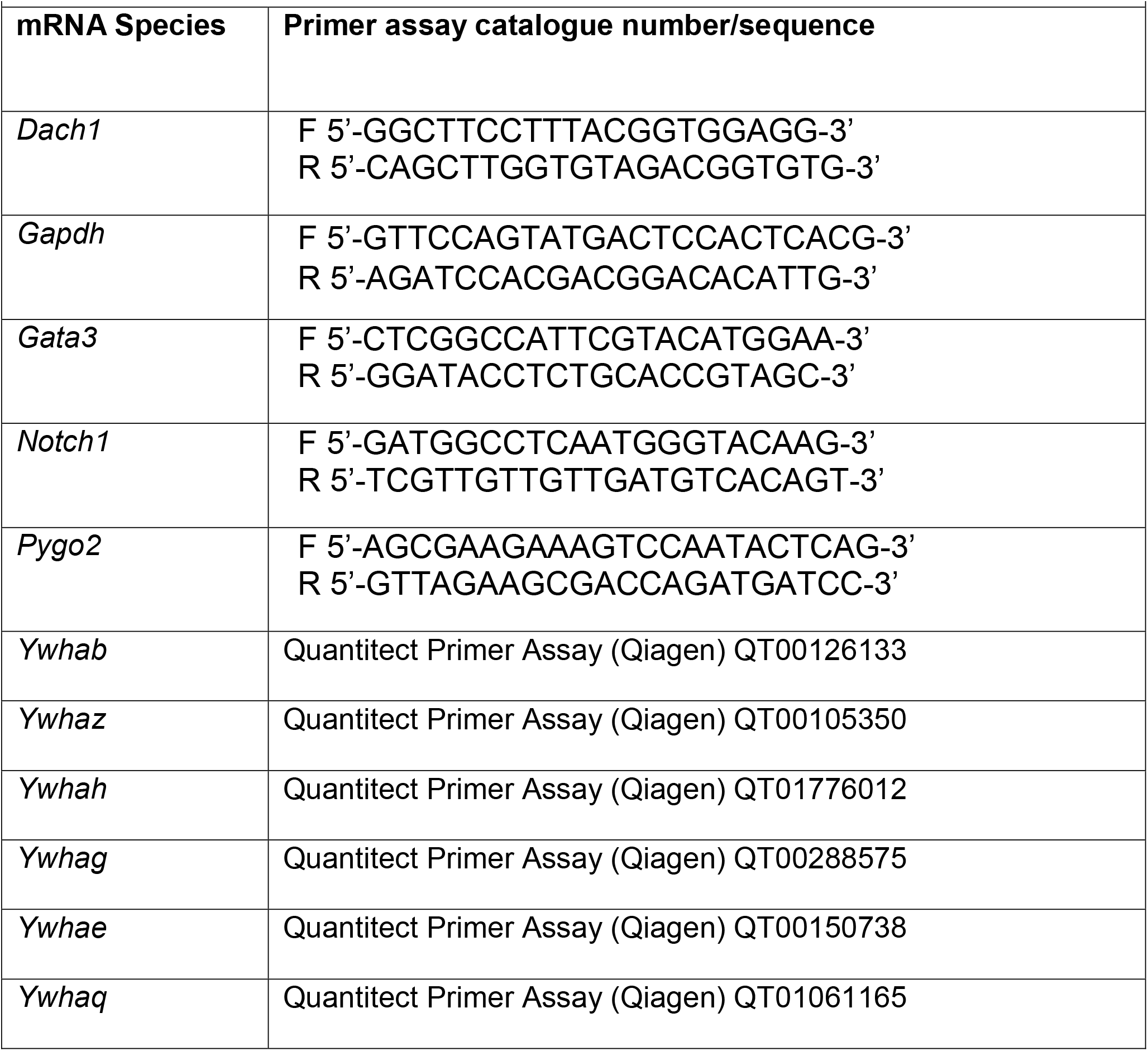
Sequences of qPCR primers and catalogue numbers of Primer assays used

Qiagen catalogue numbers and sequences for primer assays used:

### Immunohistochemistry and immunofluorescence analysis

Immunohistochemical analyses were carried out as previously described (Samuel, Lourenco, & Olson, 2011). Briefly, formalin-fixed, paraffin-embedded sections were rehydrated and immersed in 10 mM citric acid buffer at pH 6.0, boiled for 20 min, cooled, and sequentially incubated with 3% H_2_O_2_ and 5% normal goat serum in PBS. Sections were then incubated with appropriately diluted primary antibody (see Table 2), followed by Envision + System-HRP labeled Polymer (Dako). Immunolabeling was visualized with Liquid DAB + Substrate (Dako) and sections were counter-stained with hematoxylin, cleared and mounted in DEPEX (Sigma-Aldrich).

**Table 2:** Details of antibodies

| **Table 2: Details of antibodies** |  |  |  |
| --- | --- | --- | --- |
| **Antibody** | **Part No.** | **Supplier** | **Dilution** |
| 14-3-3ζ (C16) | sc-1019 | Santa-Cruz Biotechnology | 1:1000 |
| Ki-67 | VPK452 | Vector Laboratories | 1:50 |
| Cytokeratin 8 | ab53280 | Abcam | 1:200 |
| Cytokeratin 14 | NCL-LL002 | Leica Microsystems | 1:100 |

Immunofluorescence analysis was carried out as for immunohistochemistry, eliminating incubation with H_2_O_2_. Horseradish peroxidase-labeled secondary antibodies were replaced by AlexaFluor 488 or 594 conjugated secondary antibodies (Thermo Fisher Scientific) and sections were mounted in VECTASHIELD mounting medium containing DAPI (Vector Labs).

Immunohistochemically labeled sections were scanned using a NanoZoomer S60 Digital Pathology (NDP) System (Hamamatsu Photonics). Images were quantified using the Positive Cell Detection algorithm in QuPath Software (Bankhead et al., 2017).

Immunofluorescence images were acquired using an LSM 700 confocal laser scanning system (Zeiss). ImageJ was used to calculate integrated density (intensity) by signal per region of interest (duct) within each image, after conversion to a binary image based upon a single manually determined threshold value applied across all images as previously described (Kular et al., 2015).

### Terminal deoxynucleotidyl transferase dUTP nick end labeling (TUNEL)

Apoptotic cell analysis was performed using an in situ Cell death detection Fluorescein kit (Roche). Formalin-fixed, paraffin-embedded sections were rehydrated treated according to manufacturer’s instructions protocol for difficult tissues. Briefly, slides were heated in a microwave oven at 750W for 1 minute in 100 mM Citrate buffer (pH=6.0), blocked for 30 minutes in TUNEL blocking buffer (0.1M Tris-HCl, pH 7.5, containing 3% BSA and 20% normal bovine serum), incubated with TUNEL reagent and mounted in VECTASHIELD mounting medium (Vector Labs). The ImageJ particle analysis algorithm was used to enumerate TUNEL positive nuclei.

Primary antibodies used:

## Statistical analysis

Statistical analysis was performed in GraphPad Prism software (La Jolla, CA). All figure legends indicate the statistical analysis undertaken and exact P-values calculated.

## Acknowledgements

Funding was from NHMRC (M.S.S., M.K., A.F.L., H.S.R), Cancer Council SA and the Health Services Charitable Gifts Board, South Australia. V.P was supported by Research Training Program Stipend and The Commonwealth Scholarship Program for South Australia. We would like to thank the staff of the SA Pathology Animal Care Facility and the Core Animal Facility staff at the University of South Australia. The authors thank donors to the Health Services Charitable Gifts Board of SA and the Australian Cancer Research Foundation (Cancer Discovery Accelerator) for funding the imaging equipment used.

